# A Notch-regulated proliferative stem cell zone in the developing spinal cord is an ancestral vertebrate trait

**DOI:** 10.1101/298877

**Authors:** Ricardo Lara-Ramirez, Carlos Pérez-González, Chiara Anselmi, Cedric Patthey, Sebastian M. Shimeld

## Abstract

Vertebrates have evolved the most sophisticated nervous systems we know. These differ from the nervous systems of invertebrates in several ways, including the evolution of new cell types, and the emergence and elaboration of patterning mechanisms to organise cells in time and space. Vertebrates also generally have many more cells in their central nervous systems than invertebrates, and an increase in neural cell number may have contributed to the sophisticated anatomy of the brain and spinal cord. Here we study how increased cell number evolved in the vertebrate central nervous system, investigating the regulation of cell proliferation in lampreys as basally-diverging vertebrate, and focusing on the spinal cord because of its relatively simple anatomy. Markers of proliferating cells show that a medial proliferative progenitor zone is found throughout the lamprey spinal cord. We show that inhibition of Notch signalling disrupts the maintenance of this proliferative zone. When Notch signalling is blocked progenitor cells differentiate precociously, the proliferative medial zone is lost, and differentiation markers activate throughout the medial-lateral axis of the spinal cord. Comparison to other chordates suggests that the emergence of a persistent Notch-regulated proliferative progenitor zone in the medial spinal cord of vertebrate ancestors was a critical step for the evolution of the vertebrate spinal cord and its complexity.

**Summary statement:** Vertebrates develop nervous systems with numerous cells. Study of cell proliferation in the lamprey nervous system links this to a medial proliferation zone regulated by Notch signalling, a vertebrate innovation.

## Introduction

The vertebrate spinal cord develops a precise pattern of neurons and glia under the control of multiple signalling pathways. Across the dorsal-ventral (DV) axis, ventral Hedgehog signalling and dorsal Bmp and Wnt signalling coordinate the formation of different neural populations (Gouti et al., 2015; Le Dreau and Marti, 2012). Differentiation along the anterior-posterior axis is regulated by a balance between anterior Retinoic Acid (RA) signalling and posterior FGF signalling (Diez del Corral et al., 2003). As the embryo elongates, the interface between these signals moves posteriorly, leading to a wave of cell differentiation along the spinal cord. Across the medial-lateral (ML) axis, cells close to the lumen remain in a proliferative progenitor state, forming the ventricular zone of the spinal cord. Other cells migrate laterally and differentiate (Gouti et al., 2015). Spinal cord cells thus integrate information across all three spatial axes for appropriate position-specific differentiation.

The Notch signalling pathway regulates numerous aspects of vertebrate development, and in the spinal cord it regulates neurogenesis by maintaining medial cells in a proliferative progenitor (stem cell) state (Myat et al., 1996). Notch signalling relies on cell-cell contact and is activated by binding of the transmembrane proteins Delta and Serrate/Jagged to Notch receptors on an adjacent cell. Upon ligand binding, Notch suffers two proteolytic cleavages; the first is catalysed by ADAM-family metalloproteases, while the second is carried out by the □-secretase enzyme complex (Bray, 2006; Kopan and Ilagan, 2009). This releases the Notch intracellular domain (NICD), which translocates to the nucleus to promote transcription of target genes in combination with the transcription factor CSL (Bray, 2006; Fischer and Gessler, 2007). The best known direct targets of NICD/CSL are the basic Helix Loop Helix (bHLH) transcription factors of the *Hes* family, which exert an inhibitory role on neuronal differentiation and act as antagonists to neural differentiation-promoting bHLH genes including the *neurogenin*, *atonal*, *ASCL* and *COE* families. Experimental manipulation of Notch signalling in mice, *Xenopus* and zebrafish has shown that loss of Notch signalling causes cells to differentiate prematurely, depleting the progenitor pool (Appel et al., 2001; Wettstein et al., 1997). Thus, continued Notch signalling seems necessary to keep a progenitor pool over developmental time, and hence for spinal cords to develop their characteristic large number of cells. These progenitor cell populations may also be the source of adult neural stem cell populations (reviewed by (Grandel and Brand, 2013)).

A long-standing question in evolutionary biology is explaining how the complex central nervous system (CNS) of vertebrates evolved. Many studies have approached this by asking how neural patterning is regulated in vertebrates and in their nearest invertebrate relatives, cephalochordates (amphioxus) and tunicates (e.g. (Albuixech-Crespo et al., 2017a; Albuixech-Crespo et al., 2017b; Holland et al., 2013)). These studies have revealed patterning differences between vertebrates and these invertebrate lineages, explaining some of the complexity seen in vertebrates. However, complexity of pattern is only part of what makes vertebrates distinct, as vertebrates also have many more cells in their central nervous systems than do tunicates, cephalochordates and most other invertebrates.

Evolving more cells in an organ system could happen by several routes although given that in vertebrate model species these cells develop from a medial progenitor pool, a simple hypothesis explaining the evolution of extra vertebrate neural cells is the evolution of mechanisms to generate and/or maintain these progenitors in significant numbers over an extended period of development. A prediction of this hypothesis is that a lasting progenitor pool and its conserved molecular regulation should be absent in invertebrate chordates with simple neural tubes, but present in all vertebrates including the earliest diverging lineage, the agnathans (Shimeld and Donoghue, 2012). Lampreys and hagfishes, the only living agnathans, split from the lineage leading to jawed vertebrates before the evolution of hinged jaws and paired appendages, though their adult central nervous systems are large compared to those of cephalochordates and tunicates. Adult spinal cord anatomy in lampreys is relatively well studied, both as a model for the analysis of neural circuitry and for its capacity to regenerate following transection (Herman et al., 2018; Shifamn and Selzer, 2015). Much less is known about the embryonic and early larval development of the lamprey spinal cord. It is not known which signalling pathways govern cell patterning in this tissue and Notch signalling has not been experimentally tested, though some gene expression data indicate it might be involved, at least in the brain (Guerin et al., 2009).

To gain insight into the evolution of the vertebrate spinal cord and the involvement of Notch signalling in this event, we studied markers of neural cell proliferation and differentiation in lampreys, and whether these are regulated by Notch signalling. We show the entire lamprey spinal cord develops a medial progenitor zone that persists over an extended developmental period. We show maintenance of this progenitor zone is dependent on Notch signalling, and that compromised Notch signalling results in loss of the progenitor pool. Lost progenitors differentiate precociously. These data, when compared to data from other chordates, demonstrate that a CNS-wide proliferative medial progenitor zone evolved before the radiation of the jawed and jawless vertebrate lineages, but after their separation form invertebrate lineages. It is hence a vertebrate innovation. We also identify subtle differences in the outcome of Notch manipulation in the lamprey and jawed vertebrate lineages, suggesting additional evolutionary change following their divergence.

## Results

### A medial proliferation zone is present throughout the lamprey spinal cord

To identify proliferating cells we cloned the proliferation markers *Proliferating Cell Nuclear Antigen* (*PCNA)* and *Musashi* (*Msi)* from *Lampetra planeri* (*L. planeri*, or *Lp*). *PCNA* is a cofactor of δ-polymerase, and is known to be expressed in the brain of a different lamprey species, *Lampetra fluviatilis* (Guerin et al., 2009). *Msi* genes encode RNA binding proteins expressed in various stem cell populations, and one member of this family is also expressed in the *L. fluviatilis* brain (Guerin et al., 2009). Molecular phylogenetic analysis show *LpPCNA* groups with other vertebrate *PCNA* sequences with strong support (Figs. S1, S2). The *L. planeri Msi* sequence grouped within other chordate *Msi* sequences, and was most similar to jawed vertebrate *Msi2*, hence we name this gene *LpMsi2* (Figs. S3, S4).

We analysed the expression of *LpPCNA* and *LpMsi2* in normal embryos from Tahara (Tahara, 1988) stage 21 to stage 29 (Fig. 1, S5). At stage 21, *LpPCNA* is widely expressed in the protruding head and in the entire neural tube (Fig. 1A). Expression is maintained in the head and neural tube at stage 22 as the embryo grows, and at stages 23 and 24 expression is clearly observed in the pharyngeal region and becomes medially restricted in the spinal cord (Fig. 1B-F). Strong expression persists through stages 25 and 26, with spinal cord expression restricted to the medial-most cells (Fig 1G, H). This pattern essentially continues through stages 27-28, though brain and anterior spinal cord expression starts to weaken (data not shown). *LpMsi2* expression is similar to that of *PCNA* (Fig. S5). Thus, both proliferation marker genes are medially-expressed in the spinal cord over an extended developmental period, from stage 23 to at least stage 28, which spans about two weeks under normal developmental conditions.

**Figure 1.**
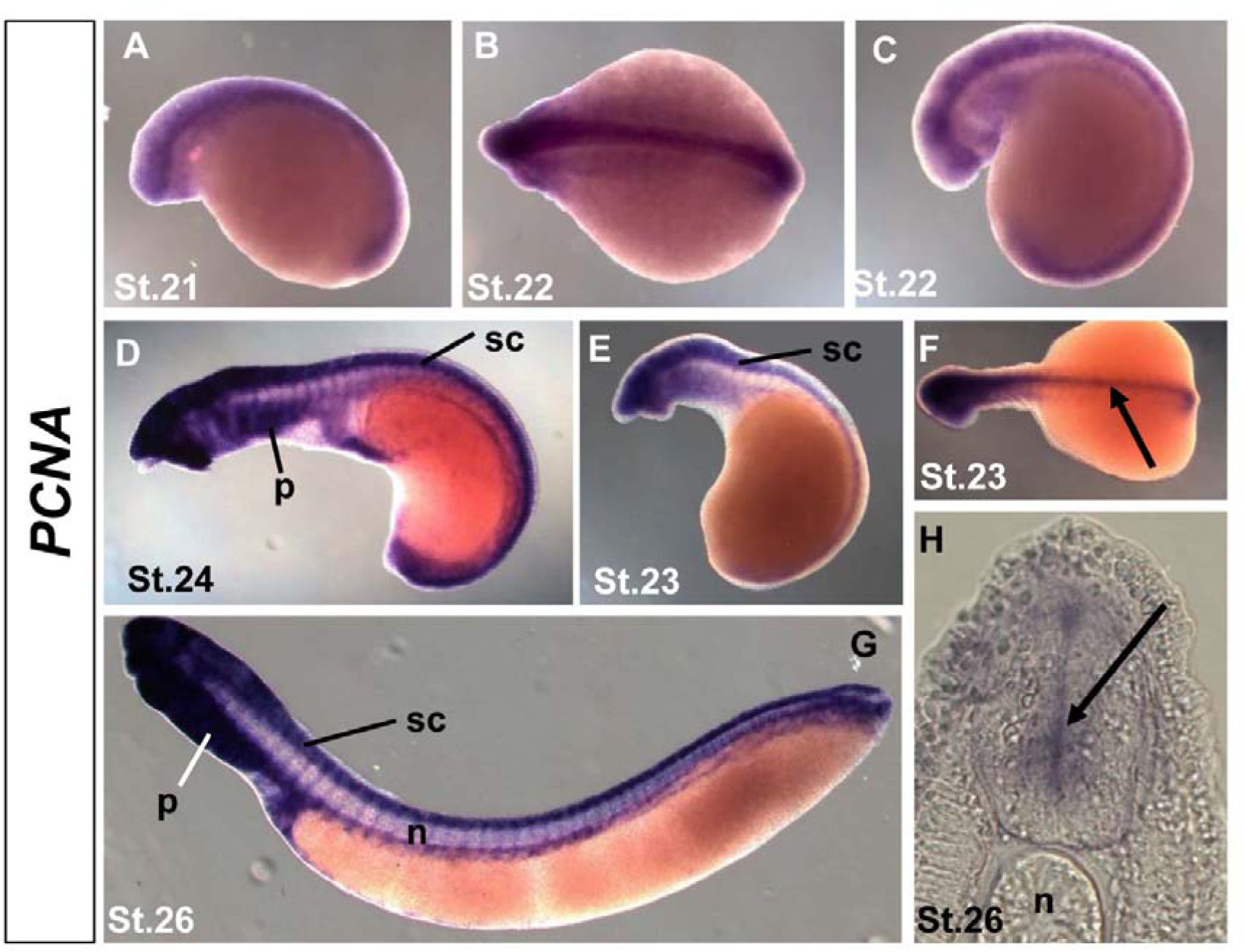
*LpPCNA* expression identifies the lamprey spinal progenitor zone. All embryos in lateral view except (B), which is a dorsal view And (H), which is a transverse section. Anterior is to the left in all images, except (H). (A) At stage 21, *LpPCNA* expression is observed along the entire neural tube and in the pharyngeal region. (B, C) At stage 22, expression is maintained in the neural tube and extends into the developing pharyngeal arches. (D-F) At stages 23 and 24, expression increases in the entire neural tube and the pharyngeal region, and in (F) can be seen to be medially restricted. At stage 26, expression increases in the entire pharyngeal region, and expression is maintained in the entire neural tube. (H) A cross-section of a stage 26 embryo through the trunk region (dorsal to the top) reveals *LpPCNA* expression in the ventricular zone (arrow). n, notochord. p, pharynx. sc, spinal cord. *LpMsi2* shows a similar pattern of expression (Fig. S5).

### Notch signalling is active and can be inhibited by DAPT treatment lamprey embryos

*Notch* ligand expression has been provisionally described in two lamprey species, and in the CNS is broadly expressed early in development before becoming more medially confined (Guerin et al., 2009; Kitt, 2013). *Hairy/Hes* genes lie immediately downstream of Notch signalling in many species (reviewed by (Iso et al., 2003)) and offer a route to understanding the strength of receipt of Notch signalling. We cloned two *L. planeri Hes* homologs (*LpHesA* and *LpHes-B*, Fig. S6, 7). *LpHes-B* expression was detected at stage 21 in the anterior CNS (Fig. 2A). Expression gradually increased and extended to most of the CNS at stages 22 and 23 (Figure 2B-D). By stage 24 it had spread through the length of the CNS (Fig. 2E, F) and maintained this pattern until stage 28 (Fig. 2G-J and data not shown). Expression was also medially restricted (Fig. 2F, G, J), as would be predicted from the location of proliferating cells.

**Figure 2.**
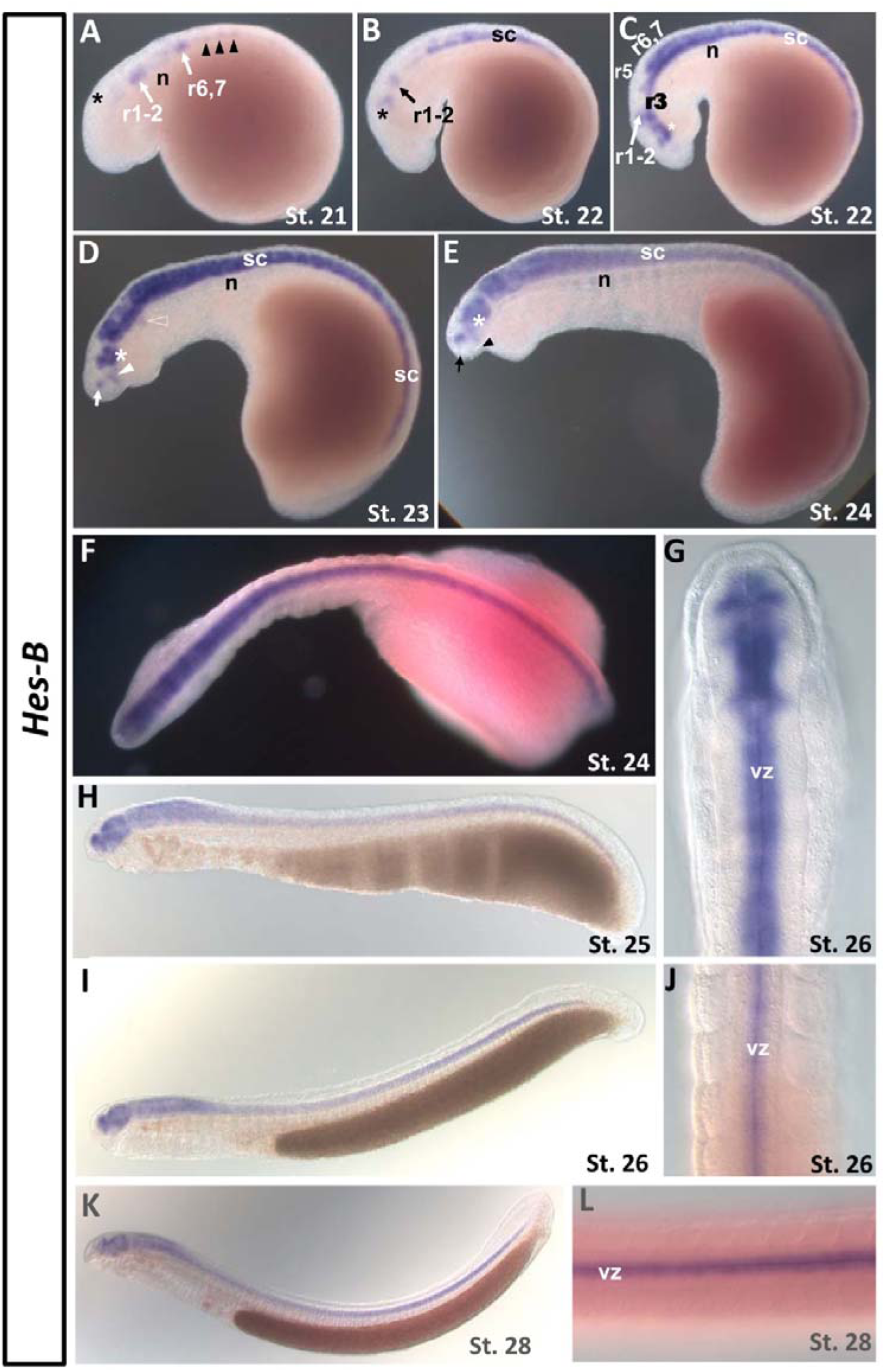
Expression of *LpHes-B* marks the progenitor zone in *L. planeri*. (A-E, H, I, K) are lateral views. (F, G, J, L) are dorsal views. Anterior is to the left in all images except in (G) and (J) in which anterior is to the top. (G) and (J) are dorsal views of the head and trunk region of a stage 26 embryo, respectively. (L) is a dorsal view of a stage 28 embryo. (A) At stage 21, *LpHes-B* is expressed in the forming neural tube. Strongest expression is seen in presumptive rhombomeres (r)1-r2 and r6-r7. Weaker expression is observed in presumptive midbrain (black asterisk) and spinal cord (black arrowheads). (B) At stage 22, expression increases in the midbrain (black asterisk) and considerably in the spinal cord. Expression is generally restricted to the ventral side of the neural tube. (C) In a slightly older stage 22 embryo, expression dramatically intensifies in the same expression domains, particularly in the midbrain (white asterisk) and spinal cord where expression extends posteriorly. Expression in r4 progressing dorsally starts to demarcate unstained r3 and r5. (D) At stage 23, expression has covered most of the rhombospinal region except in the dorsal half of rhombomeres 3 and 5 (the open arrowhead points to r4). Expression is also present in a large territory in the ventral midbrain (white asterisk). Notably the MHB is not stained. At this stage expression first appears in a discrete domain in the telencephalon (arrow) and ventral diencephalon (solid arrowhead). (E, F) At stage 24, r3 and r5 are almost completely stained while the MHB remains unstained. Expression in the midbrain extends dorsally. At this stage, all expression along the neural tube is located medially in the ventricular zone (F). (H) At stage 25, expression covers the entire neural tube except the epiphysis (G, I, J), and as also seen at stage 26 (G, I, J), is restricted to the ventricular zone. (K, L) Stage 28 embryos in lateral and dorsal views, showing expression restricted to the ventricular zone. Abbreviations: n, notochord; r, rhombomere; sc, spinal cord; vz, ventricular zone.

To further investigate this we used the Notch pathway inhibitor DAPT, which binds to and inactivates the γ-secretase complex, thus inhibiting Notch signalling by preventing release of NICD (Geling et al., 2002). This compound has been used as an inhibitor of Notch signalling in animals from chordates to cnidarians (e.g. (Lu et al., 2012; Marlow et al., 2012) and can be added over defined developmental time windows. We treated embryos with DAPT between stages 24 (when brain patterning is well advanced but the spinal cord still developing) and 26 (when the spinal cord has lengthened considerably). DAPT-treated embryos consistently bent backwards (e.g. Figure 3A, B) upon DAPT treatment, while general morphology and anatomical relationships were otherwise maintained. If DAPT inhibits Notch signalling in lamprey embryos, it should manifest in a predictable change in *Hes* gene expression, specifically *Hes* should be lost from areas where Notch signalling is active. We hence assayed *LpHes-B* gene expression in DAPT-treated and DMSO-treated control embryos. *LpHes-B* expression appeared normal in control embryos (Figure 3A, C, D). In DAPT-treated embryos, *LpHes-B* expression in the brain is similar to controls, though shows some decrease in the midbrain and dorsal hindbrain (Fig. 3, compare C and F). However expression was completely lost from the spinal cord (Fig. 3D, E, G, H). *LpHes-B* expression was also lost from the tailbud (Fig. 3G, H). We conclude from these results that DAPT effectively blocks Notch signalling in the developing lamprey spinal cord under these conditions.

**Figure 3.**
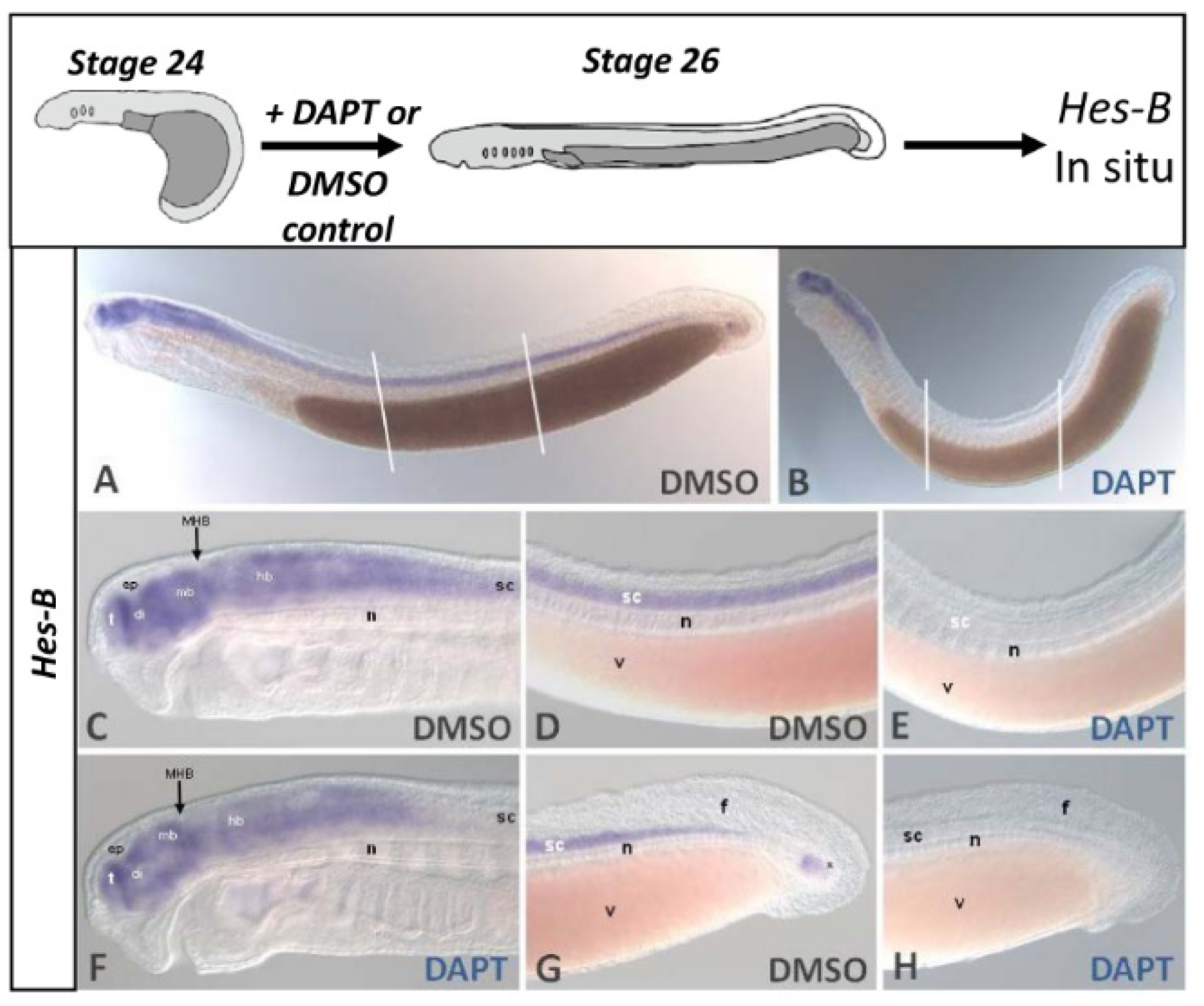
Spinal cord *LpHes-B* expression is lost when Notch signalling is inhibited. The diagrams at the top of the panel summarise the experimental design. (A, C, D, G) are control embryos. (B, E, F, H) are DAPT-treated embryos. (D, E) are corresponding regions in control and DAPT-treated embryos as indicated by lines in (A) and (B), respectively. (G) and (H) are the tail region in control and DAPT-treated embryos, respectively. In all images anterior is to the left. *LpHes-B* is expressed in the entire central nervous system and in the tail bud (asterisk in G) in control embryos. Upon DAPT treatment, expression in the spinal cord is abolished (B, E) together with expression in the tail bud (H). Note how expression in the brain is maintained overall (C, F). Abbreviations: ep: epiphysis, di: diencephalon; f: caudal fin, hb: hindbrain, mb: midbrain, MHB: midbrain-hindbrain boundary, n: notochord, sc: spinal cord, t: telencephalon, v: vitellum.

### Notch signalling blockade leads to loss of spinal cord progenitors

We investigated whether Notch signalling also maintains the proliferative state of progenitors in the lamprey spinal cord by examining the expression of *LpPCNA* in control and DAPT-treated embryos. Upon DAPT treatment, *LpPCNA* was downregulated through the posterior hindbrain and anterior spinal cord (Figure 4A, B, E, F). However LpPCNA expression was maintained in the posterior-most part of the spinal cord and tail bud region (Fig. 4A, B, G, H). In the brain, expression was reduced, although some expression was maintained in the forebrain, midbrain-hindbrain boundary (MHB), posterior hindbrain, and expression was also strong in the pharynx (Fig. 4C, D).

**Figure 4.**
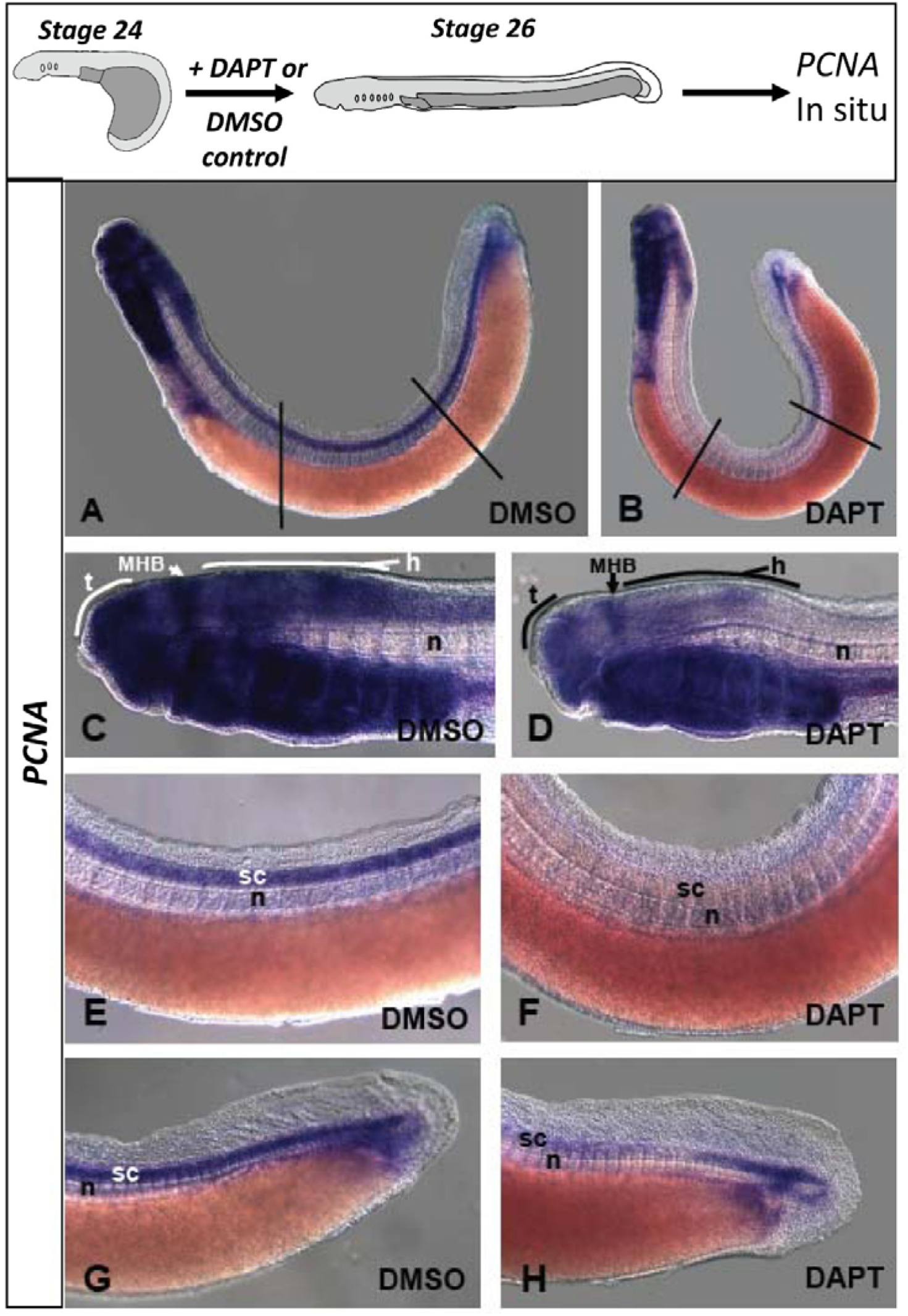
Spinal cord expression of *LpPCNA* is lost when Notch signalling is inhibited. The diagrams at the top of the panel summarise the experimental design. Control embryos (A, C, E, G) are compared to DAPT-treated embryos (B, D, F, H). All photographs show embryos with the anterior to the left. Lines in A and B show the part of the spinal cord that has been imaged in E and F. Under DAPT downregulation of PCNA is observed in the spinal cord, especially in the anterior part, and a decrease in expression in the brain is also evident. Abbreviations: t: telencephalon, MHB: midbrain-hindbrain boundary, h: hindbrain, n: notochord, sc: spinal cord.

Loss of *PCNA* expression suggests loss of proliferative progenitor cells, however it could also reflect regulation of *PCNA* expression by Notch signalling without the cell type being affected. In jawed vertebrates the progenitor zone is divided into DV zones, each formed by a specific population of proliferative cells, and each marked by well-characterised combinations of transcription factor gene expression. If progenitor cells are lost under DAPT, we reasoned the expression of these genes should also be lost. Jawed vertebrate Olig genes mark two spinal cord regions, one ventral from which motor neurons will develop, and a dorsal region spanning three progenitor zones (Alaynick et al., 2011). We first cloned a lamprey *Olig* gene which we name *LpOligA* (Fig. S6, 7). In control embryos *LpOligA* was expressed in three restricted domains of the brain (Figure 5A): two patches were observed in the diencephalon, one just above the hypothalamus and the other slightly more dorsal, both adjacent to the zona limitans intrathalamica. The third domain of expression in the brain was in the entire dorsal hindbrain. In the spinal cord two regions of expression were observed, a dorsal domain contiguous with that in the hindbrain, and a ventral domain. This mirrors the combined expression of *Olig* paralogues in spinal cords of jawed vertebrates.

**Figure 5.**
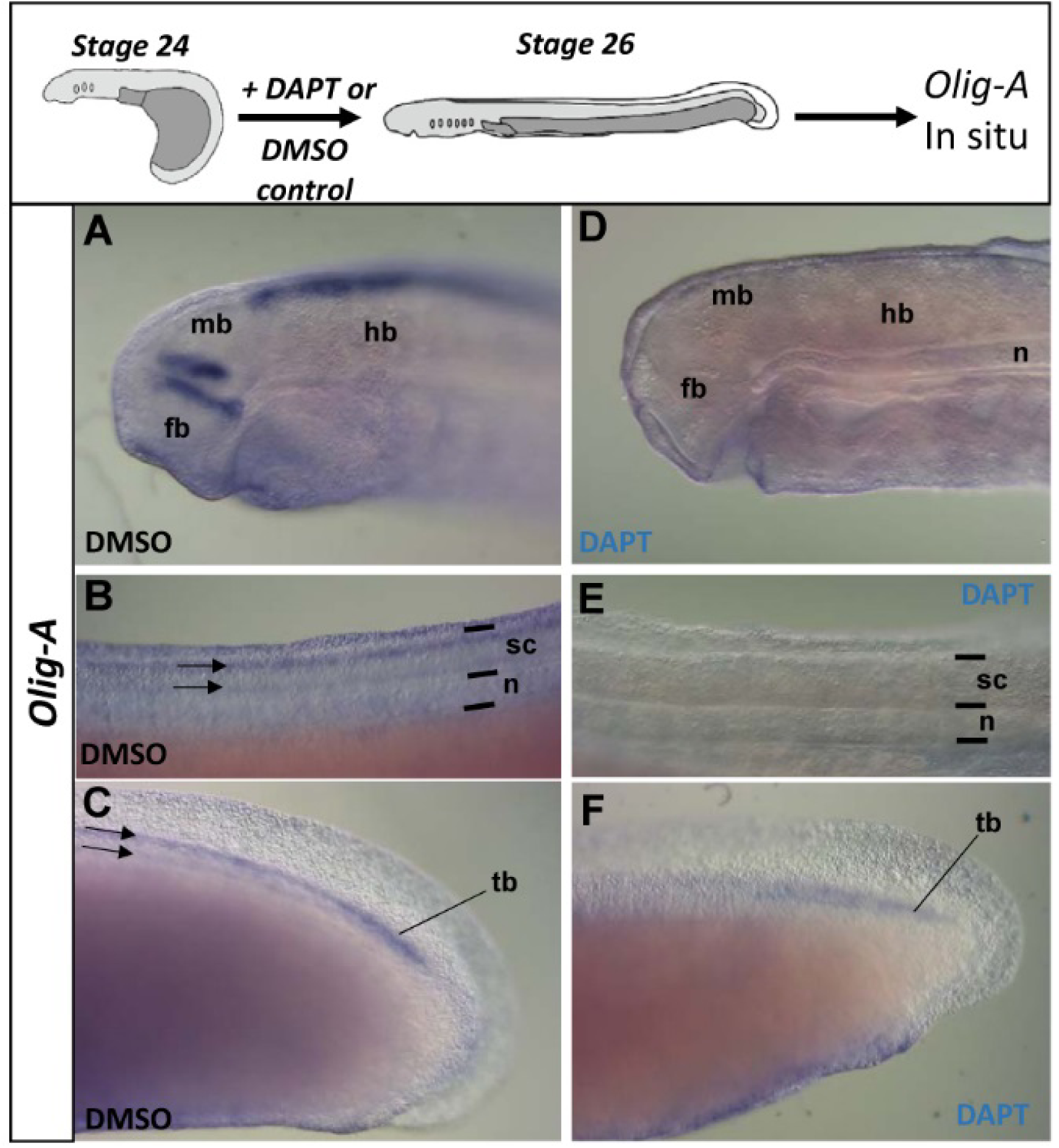
Notch inhibition blocks *LpOlig-A* gene expression. The diagrams at the top of the panel summarise the experimental design. (A-F) are lateral views, respectively of the head (A and D), dorsal trunk (B and E), and tail (C and F). Anterior is to the left in all images. (A-C) In control embryos, *LpOlig-A* expression is localised in two domains in the diencephalon, a dorsal stripe in the hindbrain, and two stripes running along the length of the spinal cord (sc; arrows on B and C). These two stripes of expression appear to merge near the tailbud (tb). (D-F) In embryos treated with DAPT, all brain and spinal cord expression is lost, though expression persists in the talibud. Black lines in (B) and (E) denote the extent of the spinal cord and notochord (n). Abbreviations: fb, forebrain; hb, hindbrain; mb, midbrain.

In DAPT-treated embryos, all three *LpOligA* expression domains in the brain were completely lost (Fig. 5D). Expression of both domains through the majority of the spinal cord was also lost, with the only remaining site a small population of cells in the very posterior, near the tail bud (Fig. 5E, F). These data support the interpretation that loss of PCNA expression in DAPT-treated embryos reflects a loss of the medial proliferative progenitor pool.

### Notch signalling blockade causes precocious differentiation in the lamprey spinal cord

To understand how the progenitor pool may have been lost, we examined the expression of the neuronal differentiation markers *LpCOE-A* and *LpCOE-B* (Fig. 6,7). *LpCOE-A* and *LpCOE-B* are broadly expressed in differentiating neurons in both CNS and PNS of normal lamprey embryos (Lara-Ramirez et al., 2017), and in the spinal cord, both genes are restricted to the more peripheral mantle layer. In DMSO-treated control embryos, both *LpCOE-A* (Fig. 6A, I, K) and *LpCOE-B* (Fig. 7A, I, K) were expressed in the peripheral region of the neural tube as in normal development. In DAPT-treated embryos, no obvious changes of expression were observed in the head (Figure 6A-D, 7A-D). However, in the spinal cord, expression of both genes expanded in two ways: first expression expanded medially, fully occupying the medial spinal cord (Figure 6J, L, 7J, L); second, expression expanded to the posterior spinal cord into areas expressing little or no *LpCOE-A* or *LpCOE-B* in control embryos (Figure 6E-H, 7E-H). These data indicate that progenitor cells are differentiating precociously both in the ventricular zone and in posterior regions of the spinal cord following Notch blockade, leading to a loss of proliferative progenitors and an increase in cells expressing differentiation markers.

**Figure 6.**
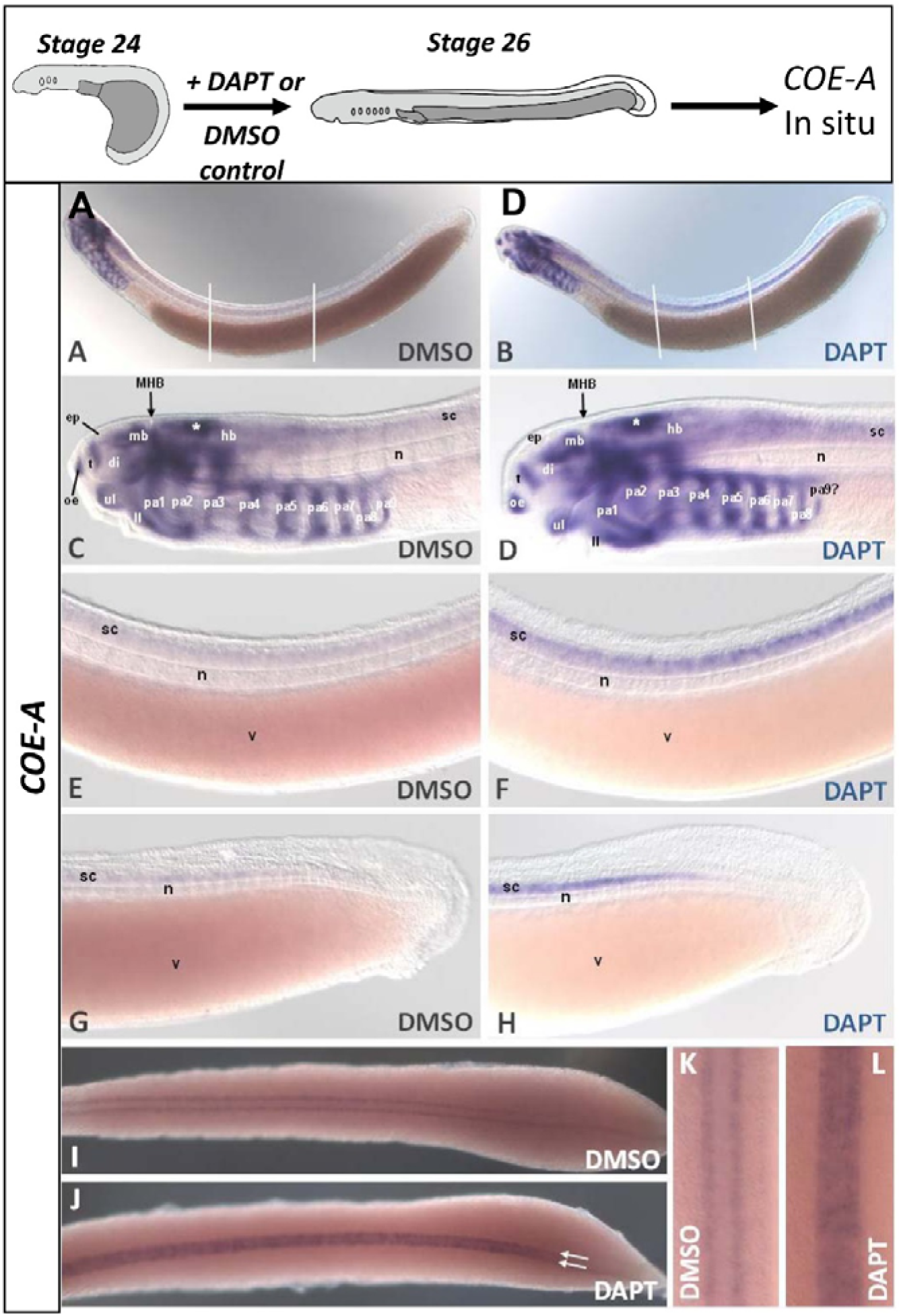
*LpCOE-A* expression expands when Notch signalling is inhibited. The diagrams at the top of the panel summarise the experimental design. (A, C, E, G, I, K) are control embryos. (B, D, F, H, J, L) are DAPT-treated embryos. (E, F) are photographs of corresponding regions in control and DAPT-treated embryos as indicated by lines in (A, B), respectively. In all images anterior is to the left except in (K) and (L) in which anterior is to the top. In the nervous system, *LpCOE-A* is expressed in the brain and faintly in the spinal cord in control embryos (A, C, E, G). Upon DAPT treatment, expression in the spinal cord it is increased as compared to control embryos (B, D, F, H). From a dorsal view, *LpCOE-A* is expressed in the spinal cord as two lateral stripes (I, K), whereas in DAPT-treated embryos expands to the middle of the spinal cord (J, L). However, in the newly-forming spinal cord at the posterior end, expression is seen as two lateral bands as in control embryos (J, arrows). Abbreviations: ep: epiphysis, di: diencephalon; hb: hindbrain; ll, lower lip; mb: midbrain, MHB: midbrain-hindbrain boundary, n: notochord; oe, olfactory epithelium; pa1-9, pharyngeal arch 1-9; sc: spinal cord, t: telencephalon; ul, upper lip; v: vitellum.

**Figure 7.**
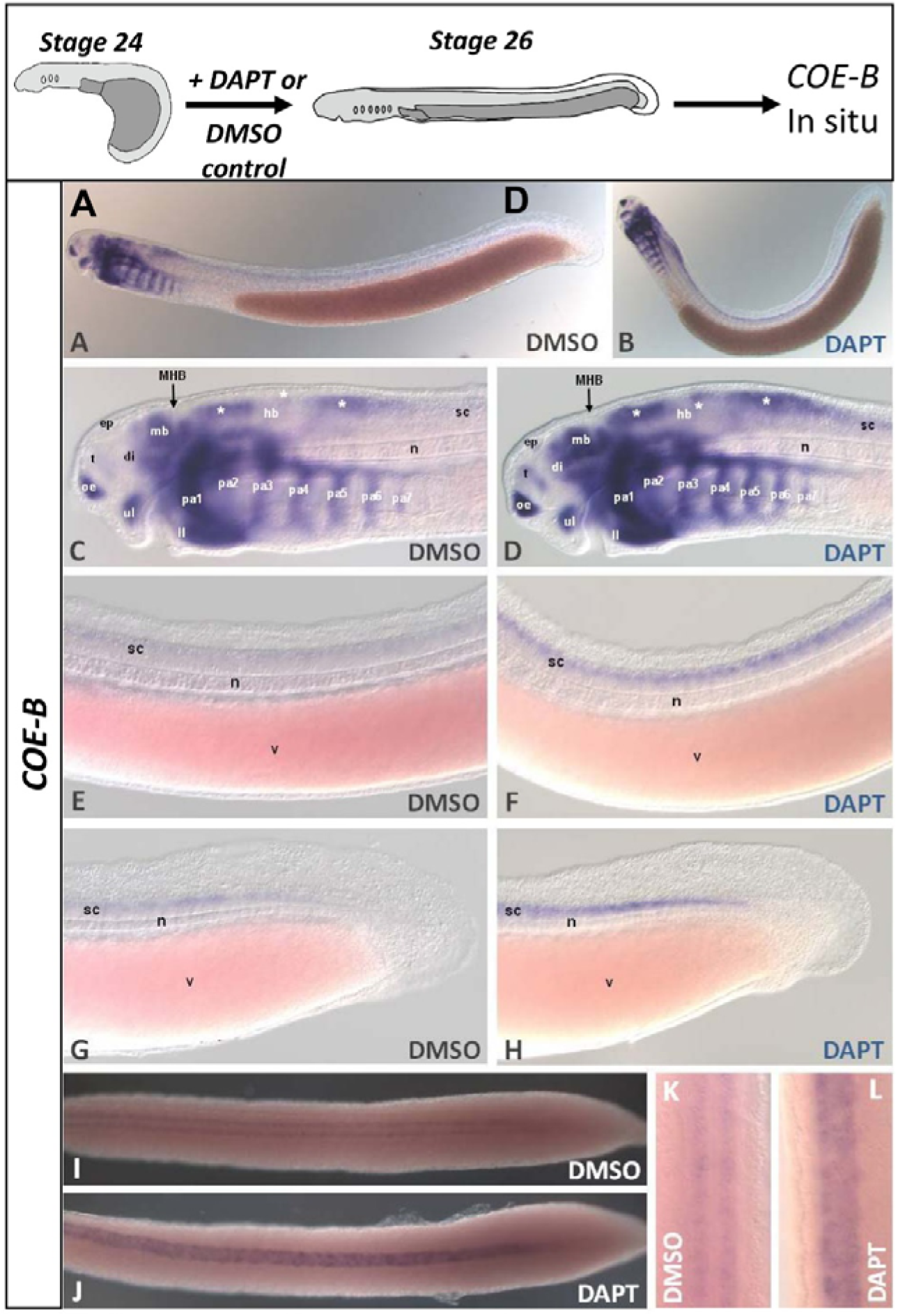
*LpCOE-B* expression expands when Notch signalling is inhibited. The diagrams at the top of the panel summarise the experimental design. (A, C, E, G, I, K) are control embryos. (B, D, F, H, J, L) are DAPT-treated embryos. (E, F) are the corresponding regions in control and DAPT-treated embryos as indicated by lines in (A) and (B), respectively. In all images anterior is to the left except in (K) and (L) in which anterior is to the top. In the nervous system, *LpCOE-B* expression is observed in restricted regions of the brain and cranial ganglia, and faintly in the spinal cord (A, C, E, G). Expression in the spinal cord is biased towards the dorsal side. Upon DAPT treatment, expression in in the spinal cord iis increased as compared to control embryos, though preserving its dorsal position (B, D, F, H). From a dorsal view, *LpCOE-B* in control embryos is seen in the spinal cord as two lateral stripes (I, K), whereas in DAPT-treated embryos expression expands into the middle of the spinal cord (J, L). Abbreviations: ep: epiphysis, di: diencephalon; hb: hindbrain; ll, lower lip; mb: midbrain, MHB: midbrain-hindbrain boundary, n: notochord; oe, olfactory epithelium; pa1-7, pharyngeal arch 1-7; sc: spinal cord, t: telencephalon; ul, upper lip; v: vitellum.

### Blockade of Notch signalling alters neural patterning

The reduction of proliferative cells in the ventricular zone and their concomitant premature differentiation could indicate a simple “speed-up” in the differentiation process while maintaining a normal distribution of cells. Alternatively, it could also result in an alteration of cell patterning. We noted that the different spatial distributions of *LpCOE-A* and *LpCOE-B* transcripts along the DV axis was generally maintained in DAPT-treated embryos despite their medial expansion (Fig. 6E, F and Fig. 7E, F). This preservation was most clear with *LpCOE-B*, which has a dorsally-biased expression that was maintained upon DAPT treatment (Figure 7E, F). However, increased expression of both *COE* genes in the anterior spinal cord also adopted a patchy pattern, suggesting the formation of clusters of differentiated cells (Fig. 6F, J, Fig. 7F, J). We reasoned that this could indicate DAPT was interfering with lateral inhibition regulated by notch, and to gain further insight into this we examined *LpNgnA*, which is expressed in the ventricular spinal cord in *L. planeri* (Lara-Ramirez et al., 2015). Expression of *LpNgnA* in DMSO-control embryos was as seen in wild type embryos, that is localised in regions of the brain, the cranial ganglia, and spinal cord (Lara-Ramirez et al., 2015) (Fig. 8A, C). In DAPT-treated embryos, expression appeared normal in the head (Fig. 8B, F); however, in the spinal cord, expression resolved into a series of discrete and widely-spaced patches (Fig. 8B, E, H, I, K). These patches were absent in the anterior-most spinal cord but were progressively closer together towards the posterior end (Fig. 8E, H, I, K). These data suggest that, as well as differentiating precociously, local patterning of cells is also being affected.

**Figure 8.**
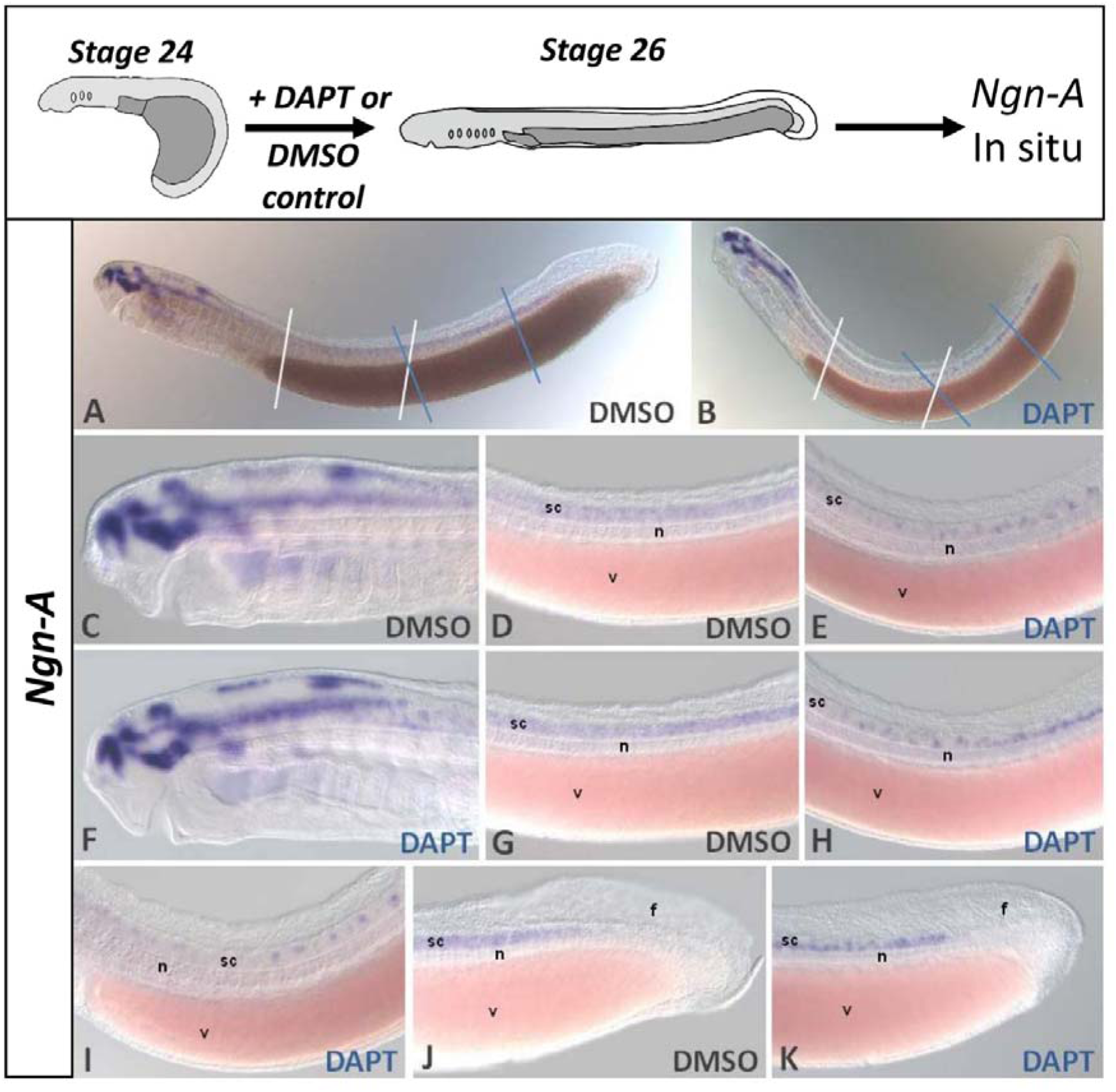
Expression of *LpNgnA* in DAPT-treated embryos. The diagrams at the top of the panel summarise the experimental design. (A, C, D, G, J) are control embryos. (B, E, F, H, I, K) are DAPT-treated embryos. (D, E) are photographs of corresponding regions in control and DAPT-treated embryos as indicated by white lines in (A) and (B), respectively. (G, H) are photographs of corresponding regions in control and DAPT-treated embryos as indicated by blue lines in (A) and (B), respectively. (I) is a photograph of a different embryo; the region of the trunk shown in this photograph overlaps with but is anterior to that delimited by white lines in (A) and (B). In all images anterior is to the left. *LpNgnA* is expressed in restricted regions of the brain in control embryos as in normal embryos (A, C; (Lara-Ramirez et al., 2015)). In the spinal cord, *LpNgnA* is expressed relatively homogenously. Under DAPT treatment, spinal cord *LpNgnA* expression changes to clusters of cells all along the spinal cord (B, E, H, I). These clusters present an irregular organisation, being bigger towards the anterior and smaller towards the posterior. Additionally, they are more densely packed towards the posterior and terminate just before the end of the spinal cord (K). Abbreviations: f: caudal fin, n: notochord, sc: spinal cord, v: vitellum.

## Discussion

The control of spatial and temporal patterning of the spinal cord of vertebrate model systems is relatively well-understood, with RA, FGF, Bmp, Wnt and Hh signals providing AP and DV axial information, and Notch signalling participating in the development of the spinal cord by maintaining a medial proliferative zone of neural precursors (reviewed by (Briscoe and Novitch, 2008)). Less well-known is when and how such patterning and its resultant complexities evolved. In this study we show that the Notch signalling is active in the spinal cord of a basally-diverging vertebrate, where it regulates proliferation and differentiation. This identifies the Notch-dependent proliferative progenitor zone as a characteristic of vertebrates, and we propose this is an important evolutionary difference to other chordates.

### The lamprey spinal cord has a proliferative ventricular zone regulated by Notch signalling

Previous studies with antibodies and RNA probes have suggested a layer of proliferating cells may lie next to the lumen of the lamprey brain (Guerin et al., 2009; Villar-Cheda et al., 2006), although how they are regulated has not been determined. Our analysis of the expression of *LpPCNA*, *LpMsi2* and *LpHes-B* corroborate this for the brain, and in addition portray a ventricular zone of proliferating cells in the spinal cord. This shows that lampreys maintain a medial progenitor zone of proliferative, undifferentiated cells throughout the developing CNS. This lasts for an extensive period of spinal cord development, more than two weeks under normal developmental conditions (Tahara, 1988). In addition, expression of the neural differentiation markers *LpCOE-A* and *LpCOE-B* marks a complementary mantle layer of differentiating cells along the entire neural tube. These data show the lamprey spinal cord resembles that of jawed vertebrates with respect to the relative placement of proliferative and differentiated cells. In particular, over a relatively long developmental time, lampreys maintain a medial proliferative stem cell zone, that is a large population of cells filling the medial region of the spinal cord from post-neurulation stages until at least approaching the point the animal becomes a fully-formed, free-living and feeding organism. This is a fundamental difference to neural development in invertebrate chordates, discussed more below.

Given the presence of a medial progenitor cell population in the lamprey spinal cord, previously reported *Notch* gene expression (Guerin et al., 2009; Kitt, 2013), and the distribution of *LpHes-B* expression (Fig. 2), we reasoned the lamprey progenitor zone may be Notch regulated. To test this we turned to DAPT, since very early neural expression of *Notch* (Sauka-Spengler et al., 2007) precludes morpholino or simple gene editing based approaches for examining its late developmental roles. While we cannot exclude the possibility that DAPT has other effects on development than those mediated by Notch signalling, DAPT has been widely used as an inhibitor of Notch across many animal phyla (e.g. (Lu et al., 2012; Marlow et al., 2012)), its target presenillin is highly conserved in lampreys (Fig. S8), and the down regulation of *LpHes-B* throughout the spinal cord of DAPT-treated lamprey embryos shows Notch signalling is affected.

Blocking Notch signalling in developing lamprey embryos results in the loss of proliferative medial cells, as visualised by loss of *LpHes*, *LpPCNA* and *LpOligA* expression, with a complementary upregulation of *LpCOE-A* and *LpCOE-B* with their expansion into the medial zone. This indicates premature differentiation of these cells. The loss of both dorsal and ventral spinal expression domains of a marker of specific subsets of progenitor cells, *LpOligA*, supports this interpretation. Thus, we conclude that the balancing of a medial proliferative zone against a peripheral differentiating zone, mediated by Notch signalling, is conserved between lampreys and jawed vertebrates, and hence a character of the vertebrate common ancestor.

Not all aspects of spinal cord cell proliferation appear to be Notch regulated. First, we note that, while the most posterior, tail bud associated domain of *LpHesB* expression is lost on DAPT treatment (Fig 3G, H), posterior expression of *LpOligA* (Fig. 5C, F) and *LpPCNA* (Fig. 4G, H) are maintained, and ectopic *LpCOE* expression does not extend into this region (Fig. 6H, 7H). This shows these cells are Notch-independent, and one possibility is their proliferation and differentiation state are regulated by tail bud derived signals such as FGF8, as reported for some jawed vertebrate model species (reviewed by (Diez del Corral et al., 2003)). Second, *LpNgnA* does not behave as a canonical proliferative zone gene. Its expression is not fully lost when Notch is blocked. Neither, as would be predicted from comparison to jawed vertebrates, is there a general, relatively homogenous, upregulation of expression (Geling et al., 2002; Nornes et al., 2008; Yang et al., 2006). Instead, when notch is blocked, broader expression is lost but small clusters of *LpNgnA* expressing cells emerge. These are reminiscent of the effects of blocking Notch signalling in Notch-dependent lateral inhibition systems, and of the *Neurogenin* dependent regulation of specific neuronal types in the vertebrate spinal cord (Korzh and Strahle, 2002; Nornes et al., 2008). The distribution of *Ngn*-expressing cells also bares resemblance to what is observed in normal amphioxus development, something discussed further below. Additional phenotyping with cell-type marker genes will be needed to understand exactly what these cell clusters are.

### Notch signalling, patterning and cell proliferation in the lamprey head

When comparing the effects of Notch inhibition in anterior regions of the neural tube we noticed a clear difference to the spinal cord. Blocking Notch signalling resulted in complete loss of *LpHes-B* expression from the spinal cord and from the tailbud, but not from the brain. In particular, *LpHes-B* expression in the forebrain, midbrain and anterior hindbrain did not appear much different between DAPT and control embryos, with only some minor changes in the midbrain and hindbrain observed. However there was a clear transition in the response to DAPT visible around the hindbrain-spinal cord junction at about the level of the 5^th^ pharyngeal slit (Fig. 3F). Despite this, *LpPCNA* expression was reduced in the brain (particularly in the midbrain and dorsal hindbrain) when notch was blocked, and *LpOligA* expression was completely lost from these regions.

This led us to consider whether, in lampreys, the brain and the spinal cord might respond differently to Notch signalling. A similar distinction of brain versus spinal cord sensitivity to Notch signalling has been suggested in the mouse, based on mice double-null mutants for *Presenilin-1* and *Presenilin-2*, in which *Shh* and *Nkx2.2* expression in the ventral neural tube is absent from the trunk but maintained in the head (Donoviel et al., 1999). These authors suggested that the effects of *Presenilin* loss are restricted to the ventral neural tube in the trunk region. However we also note that the domains from which *LpOligA* is lost, in the midbrain and dorsal hindbrain, also match the areas where *LpPCNA* expression is most reduced, and where *LpHesB* expression appears affected. The apparent differences between brain and spinal cord may therefore reflect differential sensitivity of proliferating cells to Notch inhibition, perhaps related to the time window of DAPT treatment and/or how actively they are dividing. Further dissection of proliferating cell localisation, proliferation rates and differentiation in the head will be needed to resolve this.

### The evolutionary origin of complexity in the vertebrate spinal cord: a hypothesis

The cephalochordates and tunicates are the closest living relatives to vertebrates, and the only invertebrates with neural tubes clearly homologous to those of vertebrates. Their neural tubes, however, are simpler and contain fewer cells than those of vertebrates. The ascidian larval central nervous system is composed of the sensory vesicle, the neck, the visceral or motor ganglion, and the tail nerve cord (Lemaire et al., 2002; Meinertzhagen et al., 2004). Based on morphology and gene expression, the tail nerve cord is considered to be the equivalent of the vertebrate spinal cord (Wada and Satoh, 2001). However, the ascidian tail nerve cord is composed only of ciliated ependymal cells, distributed in a row of ventral keel cells, left and right lateral rows, and a dorsal row of capstone cells, thus four cells in cross-section (Fig. 9) (Lemaire et al., 2002; Meinertzhagen et al., 2004; Wada and Satoh, 2001). There is no active cell division in the posterior CNS beyond early development. The amphioxus larval central nervous system is a tubular nerve cord and contains more cells than in tunicates, including neurons and glia along its length. It presents a transient anterior swelling called the cerebral vesicle (Wicht and Lacalli, 2005). In larvae, the CNS posterior to the cerebral vesicle seems to be made of a single layer of cells, surrounded by axon tracts (Lacalli and Kelly, 2002). Electron microscopy shows it may be a little more complicated in adults, though not much (Bone, 1960). Therefore, as well as the low cell number, a medial zone of neural progenitors and a complementary, peripheral zone of differentiated cells have not been described in either amphioxus or tunicates.

**Figure 9.**
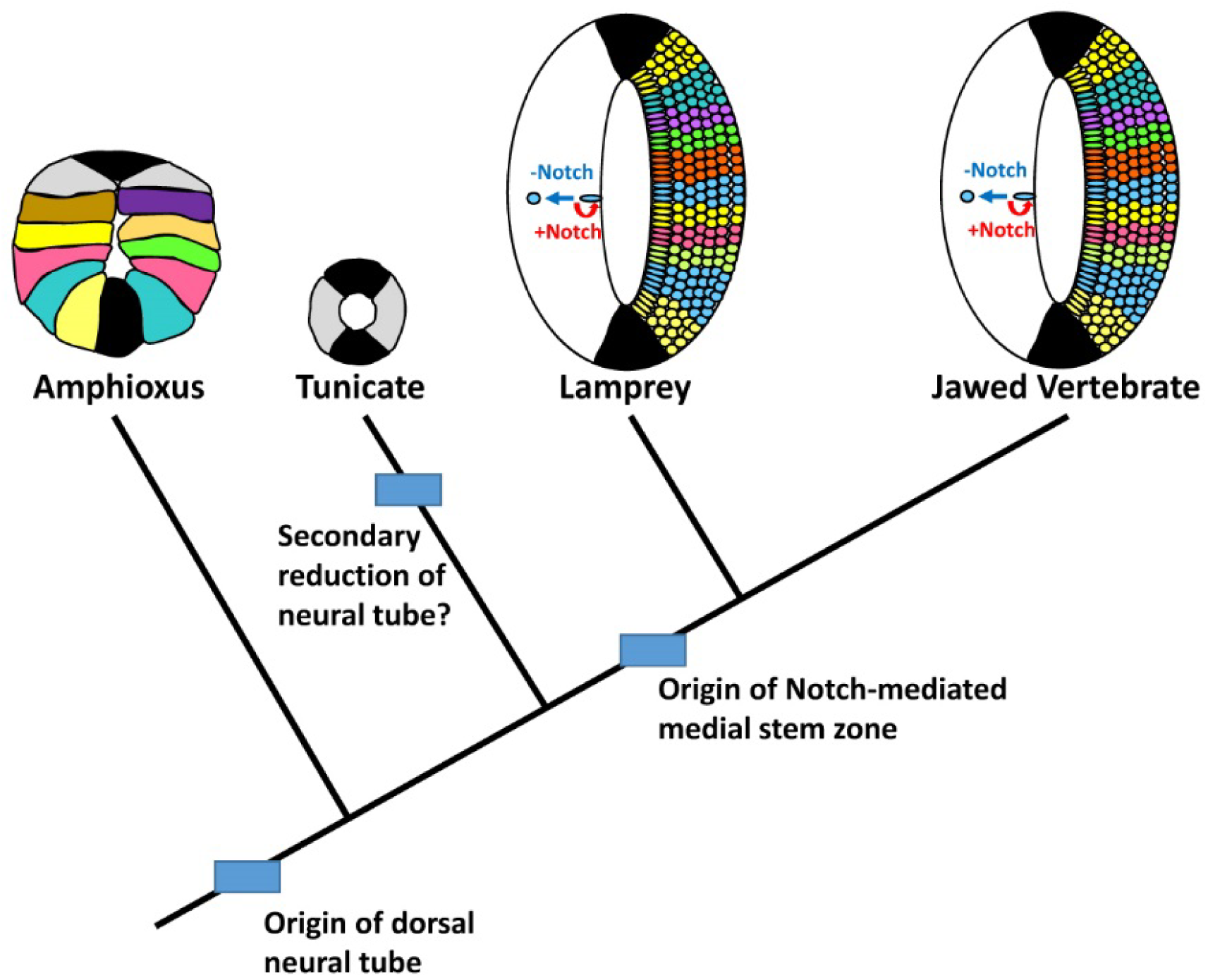
A model for spinal progenitor evolution in chordates. The diagrams at the top show schematics of the spinal cord or equivalent regions of the four major chordate lineages. The floor plate and roof plate, considered homologous between all chordates and the sources of DV patterning signals, (Corbo et al., 1997; Panopoulou et al., 1998; Shimeld, 1997; Shimeld, 1999) are show in black. In amphioxus scattered neurons of different types, some of which express marker genes found in specific subpopulations of vertebrate spinal cord cells, are found in the nerve cord posterior to the anterior swelling called the cerebral vesicle. In larvae the neural tube appears to be one cell thick (Lacalli and Kelly, 2002), though may develop more cells by the time adulthood is reached. Tunicates like *Ciona* have just 4 cells in the posterior neural tube, and no neurons, though this is inferred to be a secondary loss of complexity. Vertebrates show two key differences: (i) cells are organised into DV zones, with all cells in a zone defined by the same transcription factor gene code and (ii) a Notch-regulated stem zone is present adjacent to the lumen of the neural tube. Our data show both are also present in lampreys, and hence we conclude they are a vertebrate innovation that underlies the increase in nerve cell number and consequence neural complexity seen in vertebrates.

Expression of Notch signalling pathway components and neural HLH genes has also been analysed in ascidian tunicates and cephalochordates. In the ascidian *Ciona robusta* (formerly known as *Ciona intestinalis* type A), *Delta*, *Hes*, *Ngn* and *COE* genes are expressed in a small number of neural cells, mostly peripheral neural and sensory vesicle cells, with little or no expression in the tail nerve cord, and Notch expression persists into the nerve cord until the mid tail bud stage (Imai et al., 2004; Mazet et al., 2005; Yamada et al., 2009). In the ascidian *Halocynthia roretzi*, Notch signalling appears to be lost form the tail nerve cord by the mid tail bud stage (Hori et al., 1997). In amphioxus, the expression of *Notch*, *Delta*, *Hes*, *Ngn* and *COE* genes has been found in the neural tube including the region equivalent to the vertebrate spinal cord. *Notch* is strongly expressed in the neural tube at early neurula stages, but subsequently down regulated (Holland et al., 2001), and *Delta* expression is lost early in development from the region equivalent to the vertebrate spinal cord (Rasmussen et al., 2007). *Ngn*, *Hes,* and *COE* genes mark scattered cells in the neural tube (Beaster-Jones et al., 2008; Holland et al., 2000; Mazet et al., 2004; Minguillon et al., 2003).

Posterior CNS development in ascidians and amphioxus therefor differs from all vertebrates (including lampreys) in three key ways: (i) neither has a medial proliferative progenitor pool, (ii) in both lineages Notch signalling is maintained only through early development, and (iii) the expression of neuronal HLH genes is confined to scattered individual cells. To this we can add the observation that many of the genes that define vertebrate DV progenitor zones, and the pools of neurons that develop from them, are also only expressed in scattered individual cells in amphioxus, including members of the *Olig*, *Prdm12*, *Evx*, *Engrailed* and *Isl/Lhx* gene families (Albuixech-Crespo et al., 2017b; Beaster-Jones et al., 2008; Ferrier et al., 2001; Holland et al., 1997; Jackman and Kimmel, 2002; Thelie et al., 2015). Furthermore, when Notch signalling is blocked in lamprey development, aspects of the resultant pattern of differentiating cells resemble what is observed in amphioxus, with scattered cells rather than clearly-defined DV zones.

The vertebrate spinal cord is a far more complex structure than the equivalent in other chordates. Its development involves the production of a large number of different cell types, via a medial proliferative zone generating a peripheral zone of differentiated neurons over an extended period of development. Our data show this is present in lampreys, and hence a synapomorphic character of all living vertebrates. Thus, alongside elaboration of patterning mechanisms in early vertebrate evolution, we propose that the emergence of a Notch-regulated medial progenitor zone along the length of the CNS was a key step in vertebrate nervous system evolution (Fig. 9). In this model, a simple basal chordate nervous system is patterned across the DV axis directly into discrete cell types, marked by the expression of conserved transcription factor genes such as *Olig, Prdm12, Evx* and *Lhx*. In vertebrates two connected innovations evolve; a medial progenitor zone creates more cells, over a longer developmental time, and division of the progenitor region into DV pools forms a diversity of stem cell populations, able to form differentiated neurons over an extended developmental time. Coupled, these evolutionary innovations underlie spinal cord cell number and diversity in vertebrates.

## Materials and methods

### Animal collection and fixation

Naturally spawned *L. planeri* embryos were collected from the New Forest National Park, United Kingdom, with permission from the Forestry Commission. Fertilised eggs and embryos were collected by digging at the bottom and surrounding areas of the nests. Embryos were brought to the laboratory and placed in Petri dishes with filtered river water from the same river where they were caught. They were kept at 13-15 ºC and later fixed at different stages of development following the staging system of (Tahara, 1988). When necessary, embryos were dechorionated with fine forceps before fixation. Embryos were fixed in 4% PBS-buffered paraformaldehyde (PFA) pH 7.5, which was cooled on ice before use. Embryos were fixed in an approximately 10X excess volume of 4% PFA-PBS with respect to river water at 4 °C for at least overnight. After fixation, embryos were washed twice in DEPC-treated 1X PBS for 10 minutes each, and then dehydrated through a graded series of PBS:methanol (25%, 50%, and 75% of methanol in 1X PBS) once for 10 minutes each. Finally, they were washed twice in 100% methanol for 10 minutes each and stored in fresh methanol at −20 °C.

### Cloning and sequence analysis of *L. planeri* genes

Genes were amplified from cDNA from mixed stage *L. planeri* embryos using the following primers: PCNA: Forward 5’-GCACTCGCCAAAATGTTCGA-3’ Reverse 5’-ACCGCTGGTCTGTGAAAGTT-3’. Msi2: Forward 5’-ATTCCCCCGAAGAACACAGC-3’ Reverse 5’-GAGACGCGTAGAAGCCGTA-3’. HesB: Forward 5’-CCCCGCTGCCCACGGCAA-3’ Reverse 5’-GCTTTTTGAGACATTGGCTTTTATTGACATTC-3’. OligA: Forward 5’-GATGAAGAGCTTGGGCGGAA-3’ Reverse 5’-CTTCATCTCGTCCAGGGAGC-3’. Sequences have been deposit GenBank, accession numbers MH020217 to MH020220. Sequence analysis and manipulation were performed using MAFFT v6.864b (Katoh and Toh, 2008) The parameter for strategy for MAFFT alignments was set as “auto”. All other parameters as the defaults. Phylogenetic trees were constructed using MrBayes 3.1, using the mixed model (Ronquist and Huelsenbeck, 2003). One million generations were performed and parallel chains checked for convergence, before discarding the first 25% of trees for calculating consensus trees and posterior probabilities.

### DAPT treatments and in situ hybridisation

Live embryos were treated with DAPT (N-[N-(3,5-Difluorophenacetyl-L-alanyl)]-S-phenylglycine t-Butyl Ester), a drug that inhibits the Notch signalling pathway. After collecting, embryos were placed in Petri dishes with water from the same river where they were collected from and kept in an incubator at 13-15 °C. Embryos that reached developmental stage 24 according to (Tahara, 1988) staging classification were placed in 4-well Nunc dishes and 100 µM DAPT (from a 10 mM stock solution dissolved in DMSO) in filtered river water was added. All embryos were allowed to develop until control embryos (treated with the same volume of DMSO) reached stage 26 at 13-15 °C. Embryos were fixed in 4% PFA-PBS at 4 °C overnight, before processing for in situ hybridisation. In situ hybridisation experiments were carried out as previously described (Lara-Ramírez et al., 2014).

## Acknowledgements

We thank the Forestry Commission of England for permission to collect lamprey embryos, Dr. Jo Begbie for access to histology facilities, and Dr. Tatjiana Sauka-Spengler for access to *P. marinus* sequence and Notch gene expression data.

